# mir-218 modulates ARF6 expression and inhibits pancreatic ductal adenocarcinoma invasion

**DOI:** 10.1101/2020.09.29.319236

**Authors:** Brenna A. Rheinheimer, Ronald L Heimark

## Abstract

**Background:** Pancreatic ductal adenocarcinoma is an extremely malignant disease with the majority of patients having metastatic disease upon diagnosis. Recently, it was shown the SLIT2 plays a regulatory role in melanoma invasion through the control of invadopodia. Therefore, we sought to determine the mechanism behind miR-218 inhibition of pancreatic cancer invasion.

**Methods:** mir-218 target genes were discovered using three miRNA target prediction websites. Modulation of mir-218 target ARF6 was determined by transfection with antagomirs and mimics to mir-218. Pancreatic cancer migration and invasion were measured using Boyden chamber assays. Invasion was further measured using matrix degradation assays.

**Results:** We found that mir-218 modulates ARF6 RNA and protein expression in pancreatic cancer. mir-218 did not inhibit pancreatic cancer cell migration, but did inhibit invasion. mir-218 did not affect the formation of invadopodia in pancreatic cancer cells, but did inhibit the total area of matrix degradation caused by functional invadopodia.

**Conclusions:** This work suggests that miR-218 is a suppressor of pancreatic ductal adenocarcinoma invasion through a pathway that regulates invadopodia maturation.

## Introduction

Metastasis is the number one cause of patient mortality in those diagnosed with cancer. Tumor cells with the ability to migrate and invade neighboring normal tissue have the potential to seed and grow into separate lesions at secondary sites. miR-218 has been shown to have tumor suppressive properties in several cancer types^179–181,183^ including pancreatic cancer^184^ and several targets for miR-218 have been identified. In addition, SLIT2 was found to play a regulatory role in melanoma invasion through the control of invadopodia^306^ which presents the question of whether mir-218-1 functions in a similar manner since there is evidence that both the host gene and the intronic miRNA can have regulatory roles on the same pathway. Therefore, we sought to determine the mechanism behind miR-218 inhibition of pancreatic cancer invasion.

To spread within tissue, tumor cells utilize the same mechanisms that normal cells use to migrate and invade during biological processes such as immune cell trafficking and wound healing. In order for a single cell to migrate and invade, it must modify its shape and stiffness to interact with the extracellular matrix through a cycle of interdependent signaling events^307^. Initial propulsion and elongation is driven by actin polymerization^308^. Growing protrusions interact with the extracellular matrix through integrin proteins to form a focal contact^309^ allowing the cancer cell to adhere and move along its substrate through the recruitment of surface proteases near the attachment site^310,311^. Extracellular matrix degradation then occurs as the cell body advances forward likely providing the space required for cell migration and invasion. Contraction of the cell body is done through myosin II which is induced by the small G-protein RHO and its downstream effector ROCK^312,313^. Through other mechanisms yet to be defined, the cell-extracellular matrix contacts in the back of the cell then dissolve allowing the trailing end of the cell to glide forward^314^.

A cell migration event can be considered collective if 1) the cells move together making contact with one another at least some of the time and 2) if the cells affect one another while migrating. One common type of collective cell migration is sheet migration where cells maintain close contact and continuity—typically during wound healing. Epithelial and endothelial cells in a monolayer perform sheet migration if a gap is created^315^. Both cells at the front edge and those behind it are actively motile; however, cells at the front generally exhibit more protrusive activity^316^. Another form of collective cell migration is sprouting which is characterized by the formation of a multicellular outgrowth from a preexisting structure as seen in the formation of vascular structures. The outgrowth has a leading tip cell that maintains contact with the other cells in the structure^315^. Finally, branching morphogenesis is used by cells in the formation of complex 3D structures such as the ductal system throughout the mammary gland. Unlike sprouting, branching morphogeneis does not utilize a tip cell^315^.

Collective migration is also involved in the dissemination of tumor cells. Epithelial-derived tumor cells contain the ability to spread as groups or sprouts^315^. In addition, the activity of matrix metalloproteinases is critical for collective tumor cell migration^317^. Though pancreatic ductal adenocarcinoma cells have been shown to invade individually^221^ they have also been shown to migrate collectively as well. The first mechanism is through tumor budding. Tumor budding can be defined as de-differentiated small clusters of cells at the invasive front of gastrointestinal carcinomas. Using a cohort of 117 whole tissue sections of primary human pancreatic ductal adenocarcinoma patients, low-grade budding (0-10 buds across ten different images) was observed in 29.7% of cases while high-grade budding (greater than 10 buds across ten different images) was observed in 70.3% of cases^318^. In the same study, high-grade budding was linked to advanced tumor stage, lymphatic invasion, and decreased disease-free and overall survival. Transgenic expression of podoplanin—a mucin-like protein—using the Rip1Tag2 mouse model of mouse β cell carcinoma promoted invasion of β cell tumors^319^. This invasion did not require an epithelial-to-mesenchymal transition event, but did lead to the activation of the steps necessary for collective cell migration, namely: 1) filopodia formation, 2) adhesion onto the extracellular matrix, 3) activation of matrix metalloproteinases, and 4) increased cell migration^319,320^.

Matrix metalloproteinases (MMPs) are a family of zinc-dependent endopeptidases^321^ that are involved in a myriad of cellular processes including cancer^322^. MMPs are expressed in an enzymatically inactive state and becomes proteolytically active through removal of the pro-domain either intracellularly by furin or extracellularly by other MMPs or serine proteases^323^. Localization and compartmentalization of MMPs also dictate their biological function, and while some MMPs are secreted (i.e. MMP2 and MMP9), others like MMP14 are integral membrane proteins. Several MMPs interact with cell surface receptors or localize to specific areas of the extracellular matrix leading to increased MMP activity^324^. A central role in invasion and metastasis is degradation and remodeling of the extracellular matrix. Matrix remodeling by MMP14 (MT1-MMP) patterns tissue to allow for single cell as well as collective cell migration and invasion^317^.

Matrix metalloproteinases have also been shown to be involved in an early step of pancreatic cancer formation. Acinar-to-ductal metaplasia has been proposed as an initiating mechanism for pancreatic ductal adenocarcinoma formation^325,326^. Expression of MMP-7 was found in approximately 98% of well-differentiated pancreatic ductal adenocarcinoma tissue samples as well as 100% of associated metaplastic ductal epithelium in pancreatic cancer patients^327^. Crawford et al. also showed using a mouse model of chronic pancreatitis-associated acinar-to-ductal metaplasia that MMP-7 was essential for the propagation of metaplastic epithelium by regulating acinar cell apoptosis suggesting that MMP-7 activity influences tumor formation by exposing the epithelium to apoptotic selective pressure. Years later it was discovered that not only is MMP-7 required for Notch activation leading to the dedifferentiation of acinar cells, but is sufficient to induce this process^328^. MMP-9, which has been shown to be expressed at high levels in both pancreatic intraepithelial neoplasia and pancreatic cancer^329^, also increased acinar-to-ductal metaplastic events^330^ further suggesting a role for matrix metalloproteinases in the formation of pancreatic ductal adenocarcinoma. Recently, it was discovered that membrane-bound MT1-MMP plays a role in pancreatic fibrosis and desmoplasia^331,332^ by releasing matrix-associated TGF-β and is also required for pancreatic cancer invasion through the induction of epithelial-to-mesenchymal transition by CD44-induced Snail expression^333^.

Invadopodia are actin-rich pseudopod structures selectively found on the ventral surface of cancer cells that carry out an essential process of cancer cell invasion: degradation of the extracellular matrix through the matrix metalloproteinase MT1-MMP^334–336^. There are three phases in the life of an invadopodium: initiation, assembly and maturation. Initiation of invadopodia occurs when the activation of Src through growth factor receptors or focal adhesion kinase leads to the phosphorylation of TKS5. Once phosphorylated, TKS5 localizes to areas containing phosphatidylinositol-3,4-bisphosphate and cortactin^335^ leading to the activation of signaling cascades required for actin polymerization.

Invadopodia assembly requires actin polymerization where actin nucleators and polymerization activators are controlled by several signaling molecules including small GTPases, membrane phospholipids and protein phosphorylation. The Arp2/3 complex is an essential component of invadopodia with N-WASP being the major Arp2/3 complex activator^334,336^. Actin-binding proteins such as cortactin and cofilin also affect invadopodia formation where phosphorylation of cortactin releases cofilin leading to severed actin filaments creating new barbed ends for actin polymerization^337^. Once enough actin filaments have been made, dephosphorylated cortactin then inhibits cofilin’s severing activity to promote actin filament elongation and invadopodia assembly^336^.

Extracellular matrix degradation does not occur until invadopodia become fully mature and are able to secrete matrix metalloproteinases. The membrane-tethered MT1-MMP can be delivered to invadopodia in three different ways: 1) through endocytic recycling, 2) from intracellular stores via a Rab8-dependent secretory pathway, and 3) through coordinated actions of the actin cytoskeleton and the exocytic machinery^335^. Aside from its role in actin nucleation, N-WASP also plays a role in clathrin-mediated endocytosis and fission of endocytic vesicles and vesicle departure from the plasma membrane. Yu et al. showed that MT1-MMP is delivered via late endosome/lysosome trafficking to the plasma membrane of invadopodia where it is anchored by F-actin and enriched to promote degradation of the extracellular matrix in breast cancer cells^338^. WASH, another member of the WASP family of Arp2/3 activators, also plays a role in actin nucleation and the endosomal/lysosomal system. Montiero et al. discovered a general mechanism of exocytosis of MT1-MMP-positive late endosomes at the plasma membrane of invadopodia in breast cancer cells. This mechanism consisted of a two-step process of shuttling MT1-MMP to the plasma membrane: 1) WASH-dependent Arp2/3 activation controlled connections between tubular endosomal membranes to the invadopodia plasma membrane and 2) mediation of tethering MT1-MMP-positive late endosomes to the plasma membrane of invadopodia required the exocyst complex^339^. While the majority of invadopodia work has been done in breast cancer, invasion and metastasis of pancreatic ductal adenocarcinoma through invadopodia formation and maturation has also been shown^340,341^.

*ARF6* is a member of the ARF family of the Ras superfamily of small GTPases that has been shown to be involved in the acquisition of migratory and invasive phenotypes in a variety of cancers both *in vitro* and *in vivo*^342,343^. ARF6 is activated and converted to the GTP-bound form through guanine nucleotide exchange factors (GEFs) which determine the amount and location of the active form of Arf proteins^344,345^. In contrast, ARF6 is turned off with the aid of GTPase-activating proteins (GAPs) since Arfs bind GTP tightly but have a very low intrinsic rate of hydrolysis^346^. ARF6 has also been shown to localize to the plasma membrane and endosomal compartments where it regulates actin remodeling and endocytic membrane trafficking through its exocyst complex effector^347,348^. Interestingly, it was recently discovered that the guidance molecule *SLIT2* inhibits ARF6 activation leading to the stabilization of the interaction between N-cadherin and β-catenin at the plasma membrane of melanoma cells^306^. This SLIT2-mediated inhibition of ARF6 prevented melanoma invasion by decreasing invadopodia activity as evidenced by a decrease in the percent of melanoma cells containing the ability to degrade a gelatin matrix.

To further understand the potential role of miR-218 in the inhibition of pancreatic ductal adenocarcinoma invasion, overexpression of miR-218 with a miRNA precursor and knockdown of miR-218 with a miRNA antagomir was used to modulate miR-218 levels in pancreatic cancer cell lines. Knockdown of *ARF6* with siRNA was used as a positive control. To study this, one KRAS-dependent pancreatic cancer cell line (Capan-1) and one KRAS-independent pancreatic cancer cell line (Hs 766T) was chosen. Both of these cell lines express mir-218-1, but do not express its host gene *SLIT2*. The precursor to miR-218 led to a decrease in *ARF6* expression while the miR-218 antagomir led to an increase in *ARF6* expression in both pancreatic cancer cell lines. Boyden chamber assays were utilized to measure the effect of miR-218 on invasion in two-dimensions. The precursor to miR-218 led to a decrease in invasion in both cell lines while the miR-218 antagomir partially resuced the invasive phenotype of the cells. Matrix degradation assays were performed to determine the role miR-218 plays in invadopodia assembly and maturation. Modulation of miR-218 levels did not lead to a difference in functional invadopodia formation, but did affect invadopodia maturation. The precursor to miR-218 led to a decrease in fibronectin degradation in both cell lines while the miR-218 antagomir partially rescued the degradative phenotype in Capan-1 and fully rescued the degradative phenotype in Hs 766T.

## Materials and methods

### Transfection and modulation of ARF6 expression

Pancreatic cancer cell line Capan-1 was plated at a density of 3×10^4^ cells/cm^2^ while pancreatic cancer cell line Hs 766T was plated at a density of 1.5×10^4^ cells /cm^2^ in individual 6-well tissue culture plates. Each cell line was transfected for 72 hours with 100 nM Pre-miR Precursor to hsa-miR-218-5p (Ambion, Inc., Austin, TX), 100 nM Anti-miR miRNA Inhibitor to hsa-miR-218-5p (Ambion, Inc., Austin, TX), or 100 nM FlexiTube siRNA to *ARF6* (Qiagen, Valencia, CA) using 9.5 μl of Oligofectamine (Life Technologies, Benicia, CA). Total RNA was extracted using Trizol (Sigma Aldrich, St. Louis, MO) and quantified by absorption measurements at 260 nm. 1 μg of RNA was reverse transcribed using 1 μg/ml random primers and SuperScript II reverse transcriptase (Life Technologies, Benicia, CA). Primers to *ARF6* and *18s* were designed using the Roche Universal Probe Library assay design center and qPCR was performed using Quanta PerfeCTa Supermix, Low Rox (Quanta BioScience, Gaithersburg, MD) on an ABI Prism 7500 Sequence Detection System (Life Technologies, Benicia, CA). Differences in expression between transfected and control non-transfected cells were determined using the comparative Ct method described in the ABI user manual relative to *18s* for *ARF6*. Each cell line was transfected in duplicate and each RNA sample was analyzed by qPCR in triplicate. *ARF6* primers are as follows: forward primer 5’-GACTGCAAAGGCAGTATACAGGA-3’ and reverse primer 5’-CCGCTCAATTTTAGTTTTCAGAC-3’. *18s* primers are listed in Table 2.1.

### Dual-luciferase reporter assay

Dual-Luciferase reporter assays were performed according to the manufacturer’s protocol (Promega, Madison, WI). The wild-type *ARF6* 3’ UTR and the *ARF6* 3’ UTR which contains a mutation of the miR-218 binding site were separately cloned into pGL3 containing a luciferase reporter gene. Hs 766T were plated at a density of 3×10^4^ cells/cm^2^ in individual 6-well tissue culture plates. Forty-eight hours after plating, cells were transiently transfected with pGL3-ARF6, pGL3-ARF6mut or dual transfected with one of the pGL3 constructs and either *mir*Vana® miRNA Mimic miR-1 Positive Control or *mir*Vana® miRNA Mimic hsa-miR-218-5p (Thermo-Fischer Scientific, Anthem, AZ) and a control pTK Renilla reporter construct. Twenty-four hours after transfection, cells were lysed, protein concentration was determined using the BCA method, and analyzed on a luminometer. Luciferase activity was measured for 5 μg of protein as relative light units of luciferase activity to renilla activity. The light units were then normalized to the activity of pTKRenilla control and the results were expressed as fold activation over Hs 766T transfected with the wild-type *ARF6* 3’ UTR.

### SDS-PAGE and Western Blot

SDS-PAGE and western blot was performed as described in Chapter 3. The antibody to ARF6 (05-1149) was purchased from Millipore (Billerica, MA). The antibody to beta-tubulin (SC-9104) was purchased from Santa Cruz Biotechnologies (Dallas, TX).

### Invasion assays

Capan-1 and Hs 766T were transfected for 72 hours with 100 nM Pre-miR Precursor to hsa-miR-218-5p (Ambion, Inc., Austin, TX), 100 nM Anti-miR miRNA Inhibitor to hsa-miR-218-5p (Ambion, Inc., Austin, TX), or 100 nM FlexiTube siRNA to *ARF6* (Qiagen, Valencia, CA) using 9.5 μl of Oligofectamine (Life Technologies, Benicia, CA). After transfection, 5×10^4^ Capan-1 cells and 2.5×10^4^ Hs766T cells were plated in coated BD BioCoat growth factor-reduced matrigel chambers (BD Biosciences, San Jose, CA) for 24 hours. Cells were plated in serum free media in the top chamber while the bottom chamber contained complete media containing 10% FBS as a chemoattractant. After 24 hours, the chambers were stained with 50 µl of 0.5% crystal violet in 20% methanol for 1 minute, washed in distilled water, swabbed on the inside to remove noninvasive cells, and allowed to dry overnight. Images of 15 separate fields were acquired using a 40X objective on a Zeiss AxioVert 200 microscope using Axiovision Rel. 4.8 software (Zeiss, Thornwood, NY). Each cell line was transfected in duplicate and plated in triplicate. The number of invading cells per image was counted and data is presented as the mean number of invading cells. Significance was measured by a one-sided Student’s *t* test comparing each transfection treatment to the control non-transfected cell line. p-values less than 0.05 were considered significant and indicated in the figure legends.

### Matrix degradation assay

Sterile 12 mm glass coverslips were placed into individual 12-well tissue culture plates, coated with 2.5% 300-bloom gelatin/2.5% sucrose in CMF-PBS and allowed to dry overnight at 4°C. Gelatin was rehydrated with sterile water and fixed with 0.5% gluteraldehyde (Sigma Aldrich, St. Louis, MO). Fixed gelatin-coated coverslips were washed six times with CMF-PBS at room temperature and coated with 50 µg/ml HiLyte488 FITC-conjugated fibronectin (Cytoskeleton, Inc., Denver, CO) for one hour at room temperature. Coverslips were then quenched with serum free media for 30 minutes at room temperature followed by quenching with complete media. Coverslips were treated with 10 mg/ml NaBH_4_ (Sigma Aldrich, St. Louis, MO) for 15 minutes at room temperature and washed three times with CMF-PBS. Capan-1 and Hs 766T were transfected with 100 nM Pre-miR Precursor to hsa-miR-218-5p (Ambion, Inc., Austin, TX), 100 nM Anti-miR miRNA Inhibitor to hsa-miR-218 (Ambion, Inc., Austin, TX), or 100 nM FlexiTube siRNA to *ARF6* (Qiagen, Valencia, CA) for 72 hours prior to plating and then seeded at 2×10^4^ cells/well in complete media. Twenty-four hours after seeding, cells were fixed with 4% PFA, stained with 1 µg/ml Alexa Fluor 594-phalloidin (Life Technologies, Benicia, CA). Stained coverslips were mounted on glass slides using *SlowFade* Gold antifade reagent with DAPI (Life Technologies, Benicia, CA). Cells were transfected in duplicate and images of 10 fields were acquired randomly using a 100X objective on a Delta Vision Elite microscope using softWoRx version 4.0 (Applied Precision, Issaquah, WA). This represents an approximate total of 25% of cells per experimental condition. Fluorescent images were taken using the green, red, and blue channels (fibronectin, phalloidin, and DAPI, respectively). The presence of functional invadopodia was compared between each oligo transfection and control non-transfected cells. To determine the presence of functional invadopodia, the number of cells with fibronectin degradation underneath them was counted and then turned into a percentage based on the number of cells with fibronectin degradation present versus the total number of cells counted. To determine the invasive capability of functional invadopodia, fibronectin degradation and cell area were calculated using ImageJ software. Fibronectin degradation was compared between each oligo transfection and control non-transfected cells based on degradation size itself and degradation area per number of cells. Data of fibronectin degradation is shown as the size of the area degraded in pixels. For all invadopodia assays, significance was measured by a one-sided Student’s *t* test comparing each oligo transfection to the control non-transfected cell line. p-values less than 0.05 were considered significant and indicated in the figure legends.

## Results

### mir-218-1 is a modulator of ARF6

Using three independent miRNA target prediction websites (miRanda, mirDB, and Target Scan), we found that the *ARF6* 3’ UTR has the potential to be a novel target for miR-218^349–351^. Looking at the sequence of the *ARF6* 3’ UTR, the predicted miR-218 binding site is a 8-mer interaction that is conserved across species from human down to horse (Figure 4.1A). To determine whether or not miR-218 indeed binds to its predicted binding site within the *ARF6* 3’ UTR, the wild-type *ARF6* 3’ UTR or the *ARF6* 3’ UTR containing a mutated miR-218 binding site was cloned into pGL3 and transfected either alone or in combination with a miR-218 mimic. Double transfection of Hs 766T with the miR-218 mimic and pGL3 with the wild-type *ARF6* 3’ UTR led to an approximate 50% decrease in luciferase activity while double transfection of Hs 766T with the miR-218 mimic and pGL3 with the mutated *ARF6* 3’ UTR did not lead to a decrease in luciferase activity (Figure 4.1B). To determine if miR-218 modulates *ARF6* gene expression, we transfected pancreatic cancer cell lines Capan-1 and Hs 766T with a precursor to miR-218 and an antagomir to miR-218. Transfection with the miR-218 precursor led to a 40% decrease in *ARF6* expression in Capan-1 and a 60% decrease in *ARF6* expression in Hs 766T. Transfection with the miR-218 antagomir led to a 3-fold increase in *ARF6* expression over non-transfected Capan-1 cells and a 1.5-fold increase in *ARF6* expression over non-transfected Hs 766T cells (Figure 4.1C and D). Transfection of Hs 766T with the precursor to miR-218 also led to a decrease in ARF6 protein levels while transfection with the miR-218 antagomir led to an increase in ARF6 protein (Figure 4.1E). Therefore, these results indicate that *ARF6* is a novel target for miR-218 and that miR-218 is able to regulate both *ARF6* mRNA and protein levels. Paxillin—a cytoskeletal protein involved in actin-membrane attachment to the extracellular matrix and validated miR-218 target—was not found to be modulated in pancreatic cancer cell lines (data not shown).

**Figure 4.1:**
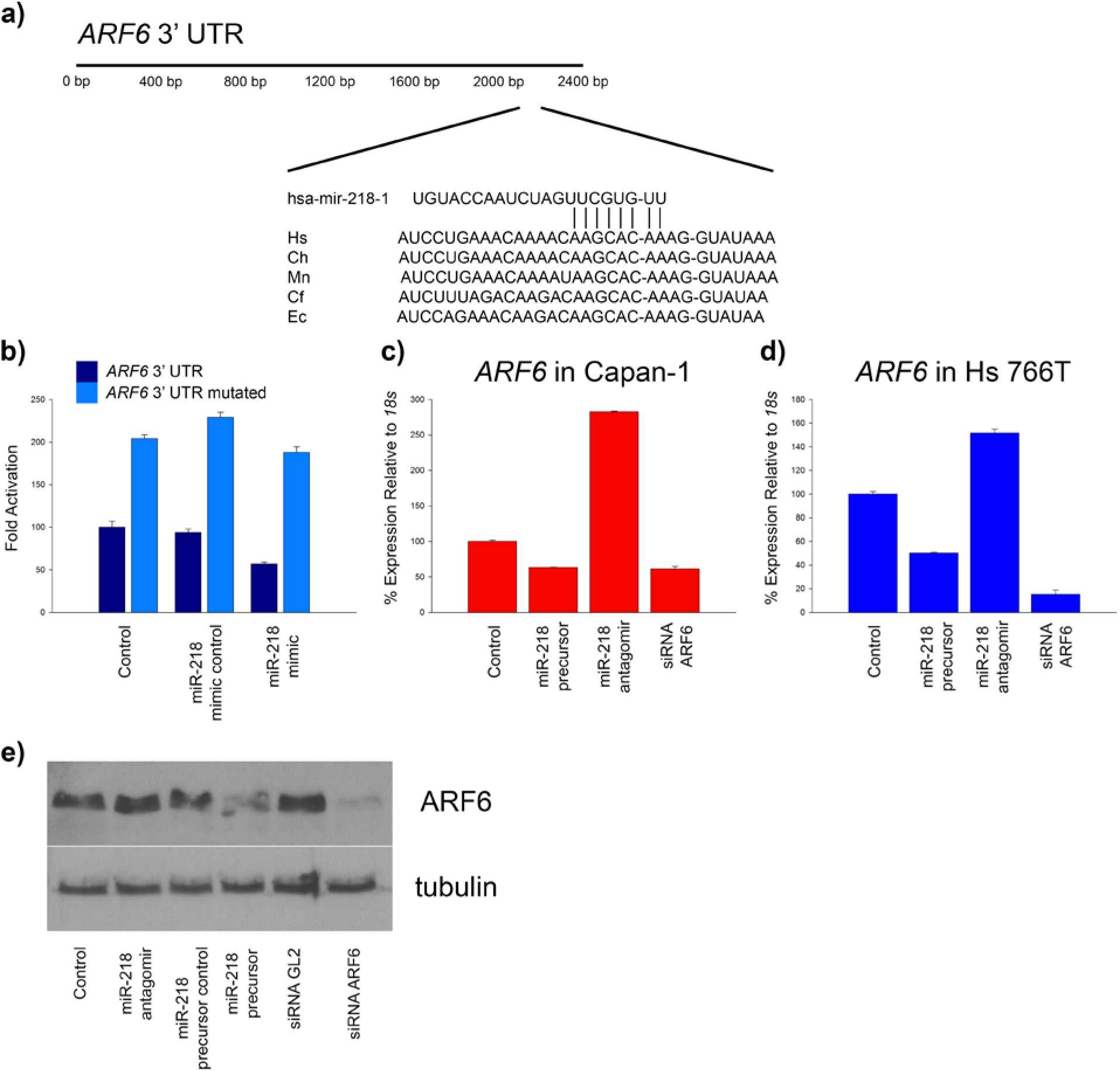
Modulation of *ARF6* expression by miR-218. (a) Conservation of the predicted 8-mer miR-218 binding site in the *ARF6* 3’ UTR. Hs = human, Ch = goat, Mn = pig-tailed macaque, Cf = dog, and Ec = horse. Fold activation of the *ARF6* 3’ UTR was analyzed using a dual-luciferase reporter assay. Hs 766T were plated at a density of 3×10^4^ cells/cm^2^ for 48 hours. Cells were then transfected with either pGL3-ARF6, pGL3-ARF6mut, or dual transfected with one of the pGL3 constructs and either a miRNA mimic control or a miR-218 mimic. Twenty-four hours after plating, cells were lysed and analyzed on a luminometer. (b) Luciferase activity showing miR-218 targeting of the *ARF6* 3’ UTR. Capan-1 and Hs 766T were transfected with 100 nM of miR-218 precursor, miR-218 antagomir, or siRNA to ARF6. Seventy-two hours after transfection, total RNA was extracted with Trizol and cDNA was prepared using random primers and Superscript II. Quantitative PCR (qPCR) was carried out and mRNA levels were normalized to control non-transfected cells relative to *18s*. Expression of *ARF6* as detected by qPCR for (c) Capan-1 or (d) Hs 766T. Additionally, after transfection, total cell lysates were extracted from Hs 766T in 2X SDS-sample buffer, protein concentration was determined, and 30 μg of protein per lane was analyzed by SDS-PAGE gel. After electrophoresis, proteins were transferred to nitrocellulose membranes and the filters were treated with a mouse monoclonal antibody to ARF6 or a rabbit polyclonal antibody to beta-tubulin. Immunoblots were developed with a chemiluminescence detection reagent. (d) Expression of ARF6 after transfection. miR-218 binds to the *ARF6* 3’ UTR and is a modulator of *ARF6* mRNA and protein expression.

### miR-218 is an inhibitor of pancreatic cancer invasion

Since pancreatic ductal adenocarcinoma is often metastatic at diagnosis, and miR-218 has been shown to play a role in inhibiting cancer cell migration and invasion^179,180,352–354^, we performed Boyden chamber assays to assess the role miR-218 plays in pancreatic cancer invasion. Transfection with both the precursor to miR-218 and siRNA to *ARF6* led to a significant (p = 0.02 and p = 9.31×10^−4^, respectively) decrease in Capan-1 invasion. Transfection of Capan-1 with a miR-218 antagomir partially rescued the invasive phenotype to 68% of non-transfected Capan-1 (Figure 4.2A and B). Transfection of Hs 766T with both the miR-218 precursor and siRNA to *ARF6* led to a significant (p = 4.69×10^−4^ and p = 0.01, respectively) decrease in invasion. Transfection of Hs 766T with the miR-218 antagomir led to a partial rescue of the invasive phenotype to 78% of non-transfected Hs 766T (Figure 4.2C and D). These results indicate that miR-218 inhibits pancreatic cancer invasion and that the suppression of the invasive phenotype does not depend on Kras since suppression of invasion occurred in cells that are both dependent and independent on Kras for growth and survival.

**Figure 4.2:**
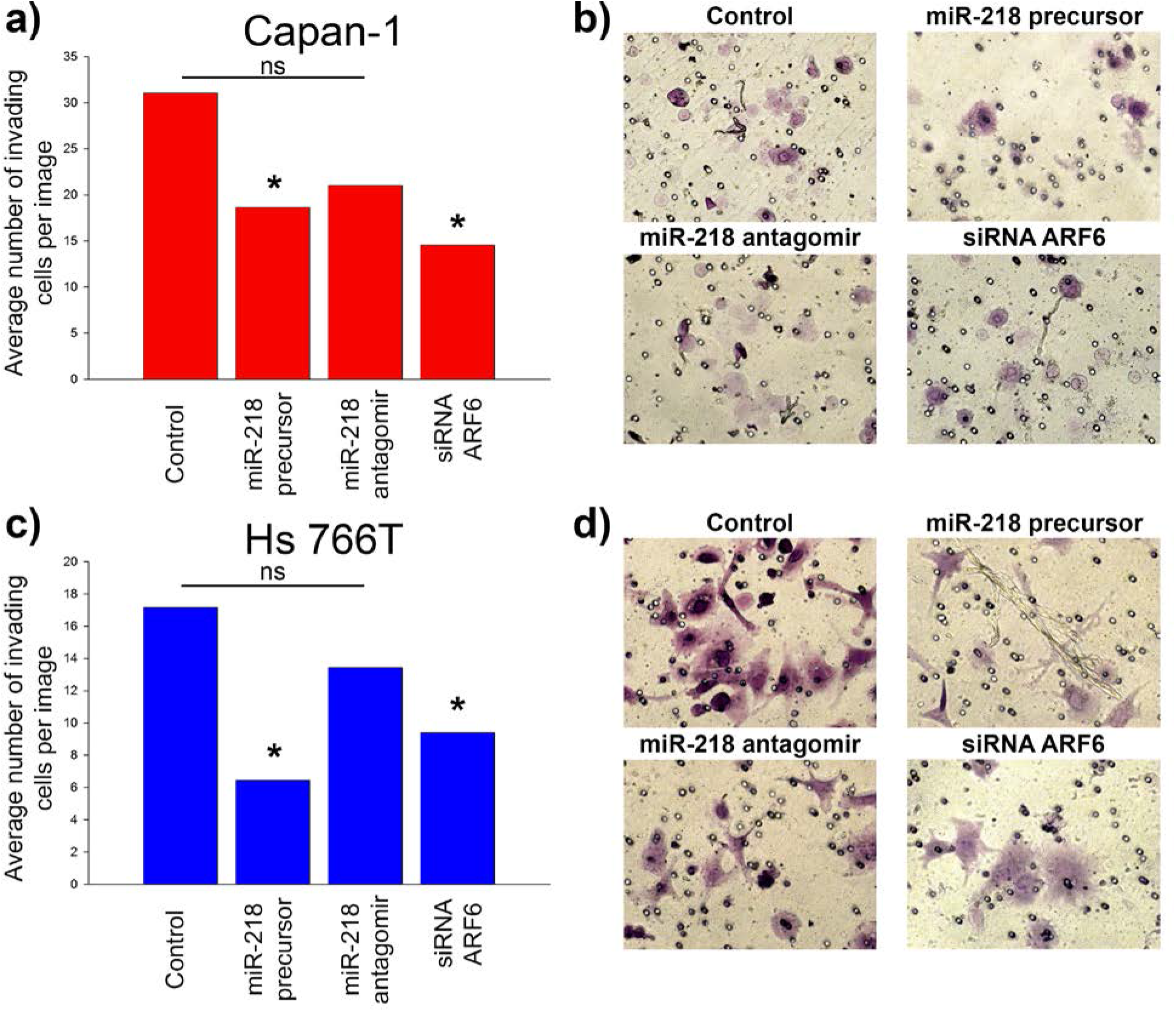
miR-218 inhibits pancreatic cancer cell invasion. Capan-1 and Hs 766T were transfected for 72 hours with 100 nM of miR-218 precursor, miR-218 antagomir, or siRNA to ARF6 prior to plating in matrigel-coated Boyden chambers. Twenty-four hours after plating, cells were stained with crystal violet. Images of fifteen separate fields were taken. Data is presented as the mean number of cells invading through the chamber for (a) Capan-1 and (c) Hs 766T. Representative images of matrigel-coated Boyden chambers stained with crystal violet for (b) Capan-1 and (d) Hs 766T. Significance was measured by a one-sided student *t* test comparing each transfection treatment to control non-transfected cells. Significance was determined as p < 0.05 and denoted with an *. ns = not significant. miR-218 inhibits pancreatic cancer cell invasion.

### miR-218 impedes extracellular matrix degradation through inhibition of invadopodia maturation, but not invadopodia formation

Since invadopodia have been shown to be involved in pancreatic ductal adenocarcinoma invasion and metastasis^340,341,355,356^, and ARF6 plays a role in invadopodia formation and maturation^342,343,357,358^, we wanted to determine if the mechanism behind miR-218 inhibition of pancreatic ductal adenocarcinoma invasion is through inhibition of functional invadopodia formation or invadopodia maturation, which was previously shown for the host gene *SLIT2*^306^. To examine this, quantitative matrix degradation assays were employed to analyze the role miR-218 plays in functional invadopodia formation and matrix degradation by invadopodia. To conclude whether or not miR-218 plays a role in functional invadopodia formation, we counted the number of cells with matrix degradation underneath them. The actin cytoskeleton of single cells was imaged using a fluorescent antibody to phalloidin at 100X using a Delta Vision Elite microscope. If matrix degradation was present in the FITC-labeled fibronectin directly underneath the cell, it was considered evidence that the cell formed functional invadopodia. Treatment with the precursor to miR-218, the antagomir to miR-218, and siRNA to *ARF6* did not lead to any difference in percentage of cells with matrix degradation underneath them in both Capan-1 and Hs 766T (Table 4.1). These results indicate that miR-218 does not play a role in inhibiting the formation of functional invadopodia.

**Table 4.1:** miR-218 does not affect the formation of functional invadopodia in pancreatic ductal adenocarcinoma cells. Sterile glass coverslips were coated with 2.5% gelatin and dried overnight. The coverslips were then rehydrated, fixed with gluteraldehyde, coated with FITC-fibronectin, quenched with media, and treated with NaBH_4_. Cells were transfected for 72 hours with 100 nM of mir-218-1 precursor, mir-218-1 antagomir, or siRNA to ARF6 prior to plating. Twenty-four hours after plating, cells were fixed with 4% PFA, stained with Alexa Fluor 594-phalloidin, and mounted with *SlowFade* reagent containing DAPI. Images of ten separate fields were taken at 100X in the red, green, and blue channels. Data is presented as a percentage of the number of cells with degradation underneath them vs. the total number of cells counted. miR-218 does not affect the formation of functional invadopodia in pancreatic ductal adenocarcinoma cells.

|  | **Control** | **mir-218-1 precursor** | **mir-218-1 antagomir** | **siRNA ARF6** |
| --- | --- | --- | --- | --- |
| Capan-1 | 35 | 25 | 30 | 30 |
| Hs 766T | 70 | 65 | 70 | 65 |

Since miR-218 does not play a role in functional invadopodia formation, I sought to determine whether or not miR-218 plays a role in the capacity of invadopodia to degrade the extracellular matrix. Transfection with the precursor to miR-218 and siRNA to *ARF6* led to a significant (p = 0.01 and p = 7.96×10^−3^, respectively) decrease in the area of fibronectin degradation in Capan-1. Treatment with the antagomir to miR-218 partially rescued the degradative phenotype to 78% of non-transfected Capan-1 (Figures 4.3A and B). The same results were seen for Hs 766T. Transfection with the miR-218 precursor and siRNA to *ARF6* led to a significant (p = 0.01 and p = 2.79×10^−3^, respectively) decrease in fibronectin degradation while transfection with the miR-218 antagomir completely rescued the degradative phenotype to non transfected Hs 766T (Figures 4.3C and D). These results indicate that miR-218 is capable of inhibiting extracellular matrix degradation by blocking invadopodia maturation and that this quality does not depend on Kras since suppression of extracellular matrix degradation occurred in cells that are both dependent and independent on Kras for growth and survival.

**Figure 4.3:**
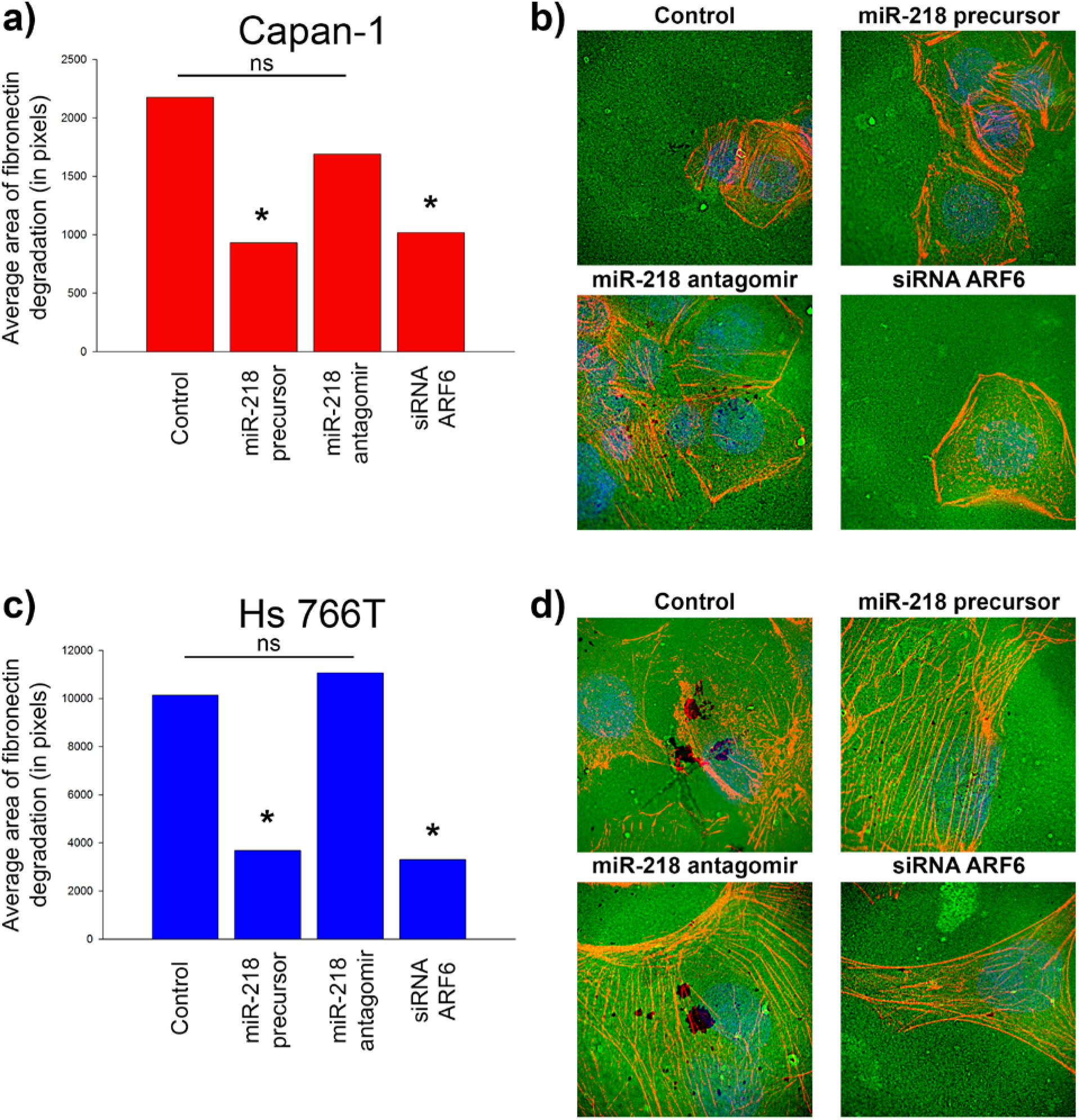
miR-218 inhibits the total area of fibronectin degradation in pancreatic ductal adenocarcinoma cells. Sterile glass coverslips were coated with 2.5% gelatin and dried overnight. The coverslips were then rehydrated, fixed with gluteraldehyde, coated with FITC-fibronectin, quenched with media, and treated with NaBH_4_. Cells were transfected for 72 hours with 100 nM of miR-218 precursor, miR-218 antagomir, or siRNA to ARF6 prior to plating. Twenty-four hours after plating, cells were fixed with 4% PFA, stained with Alexa Fluor 594-phalloidin, and mounted with *SlowFade* reagent containing DAPI. Images of ten separate fields were taken at 100X in the red, green, and blue channels. The total area of fibronectin degradation was measured in ImageJ. Data is presented as the total area of fibronectin degradation in pixels for (a) Capan-1 and (c) Hs 766T. Representative images of fibronectin degradation for (b) Capan-1 and (d) Hs 766T. Significance was measured by a one-sided student *t* test comparing each transfection to control non-transfected cells. Significance is determined as p < 0.05 and denoted with an *. ns = not significant. miR-218 inhibits the invasive capability of invadopodia in pancreatic ductal adenocarcinoma cells.

To further inspect this data, the area of fibronectin degradation was compared to the number of cells present over each area of extracellular matrix degradation. Transfection with a precursor to miR-218 and siRNA to *ARF6* led to a significant (p = 0.04 and p = 0.03, respectively) decrease in the area of fibronectin degradation per number of Capan-1 cells present while transfection with an antagomir to miR-218 led to a partial rescue of the degradative phenotype to 85% of non transfected Capan-1 (Figure 4.4A). Transfection with a miR-218 precursor and siRNA to *ARF6* also led to a significant (p = 0.02 and p = 0.01, respectively) decrease in the area of fibronectin degradation per number of Hs 766T cells present while transfection with a miR-218 antagomir led to a complete rescue of the degradative phenotype (Figure 4.4B). These results further indicate that miR-218 is capable of inhibiting extracellular matrix degradation by blocking invadopodia maturation and that this quality does not depend on Kras or the number of cells present at the site of matrix degradation.

**Figure 4.4:**
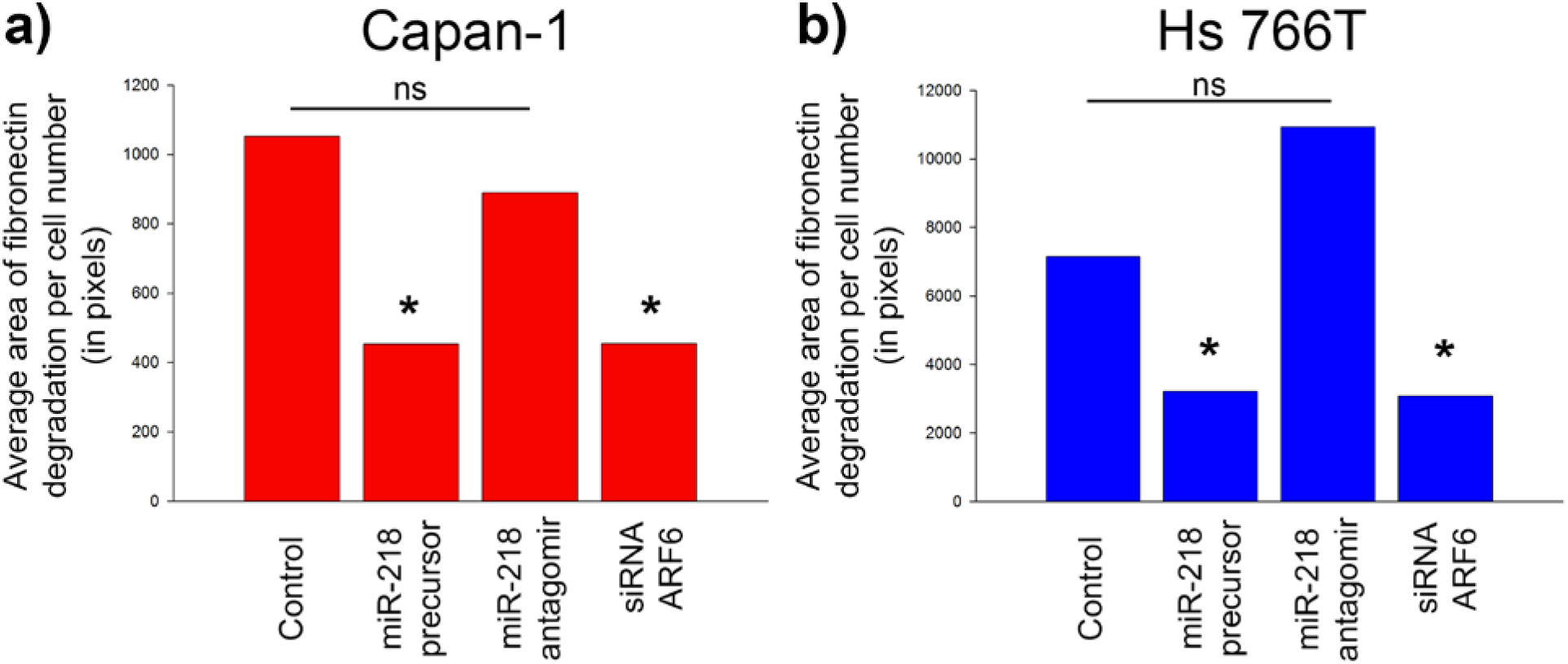
miR-218 inhibits the total area of fibronectin degradation per cell number in pancreatic ductal adenocarcinoma cells. Sterile glass coverslips were coated with 2.5% gelatin and dried overnight. The coverslips were then rehydrated, fixed with gluteraldehyde, coated with FITC-fibronectin, quenched with media, and treated with NaBH4. Cells were transfected for 72 hours with 100 nM of miR-218 precursor, miR-218 antagomir, or siRNA to ARF6 prior to plating. Twenty-four hours after plating, cells were fixed with 4% PFA, stained with Alexa Fluor 594-phalloidin, and mounted with *SlowFade* reagent containing DAPI. Images of ten separate fields were taken at 100X in the red, green, and blue channels. The total area of fibronectin degradation was measured in ImageJ and divided by the number of cells in each image. Data is presented as the total area of fibronectin degradation per cell number in pixels for (a) Capan-1 and (b) Hs 766T. Significance was measured by a one-sided student *t* test comparing each transfection to control non-transfected cells. Significance is determined as p < 0.05 and denoted with an *. ns = not significant. miR-218 inhibits the invasive capability of invadopodia in pancreatic ductal adenocarcinoma cells.

## Discussion

In the present study we have utilized overexpression of miR-218 using a miRNA precursor and knockdown using a miR-218 antagomir to determine the mechanism through which miR-218 controls invasion of pancreatic ductal adenocarcinoma cells. To begin this study, two pancreatic cancer cell lines that express mir-218-1 transcript, but do not express *SLIT2* mRNA—KRAS-dependent Capan-1 and KRAS-independent Hs 766T—were transiently transfected with either the precursor or the antagomir to determine novel targets of miR-218 known to be involved in cellular invasion. Using several different miRNA target algorithms^349–351^, the *ARF6* 3’ UTR was predicted to be a novel target of miR-218. Conservation of the predicted 8-mer miR-218 binding site within the *ARF6* 3’ UTR across species from humans down to horse suggests that this site is both functional and important in terms of regulation of *ARF6* gene expression. Double transfection of a wild-type *ARF6* 3’ UTR and a miR-218 mimic led to decreased luciferase activity while transfection of the *ARF6* 3’ UTR containing a mutation in the predicted miR-218 binding site did not. This suggests that miR-218 is capable of binding to the *ARF6* 3’ UTR to control expression. When both Capan-1 and Hs 766T were transfected with a precursor to overexpress miR-218, *ARF6* mRNA expression decreased. Conversely, when the cells were transfected with the miR-218 antagomir to knockdown miR-218, *ARF6* mRNA expression increased. The same was seen at the protein level in Hs 766T. This suggests that miR-218 has the potential to regulate cell invasion through modulation of the actin cytoskeleton and/or trafficking of matrix metalloproteinases to the plasma membrane.

ARF6 is a small G protein that is a member of the Ras family of small GTPases. It localizes to the plasma membrane and the endocytic system where it acts in a wide range of biological processes such as endocytosis, cytokinesis, and organization of the actin cytoskeleton^359^. ARF6 has been shown to activate type I PIP5K and phospholipase D^360,361^ leading to large increases in PI(4,5)P_2_ at the cell periphery. ARF6 also binds to and activates PIP5KIγ leading to the recruitment of clathrin coats in synaptic-vesicle preparations^362^ pointing to a role for ARF6 in coat-pit assembly^363^. In HeLa cells, ARF6 activity affects the recycling of integral plasma membrane proteins. There is also compelling evidence that ARF6-regulated delivery and insertion of recycling endosomal membrane at the cell surface is mediated by the vesicle-tethering exocyst complex^348^. Therefore, it is probable that ARF6 regulates the insertion of plasma membrane proteins through its GDP-GTP cycle.

*ARF6* also plays a role in actin remodeling and cell invasion. In fact, *ARF6*-mediated actin remodeling is thought to be required for the formation of pseudopods and membrane ruffles^364,365^ and cell migration^366–368^ through the activation of the Rac1 GTPase. In epithelial cells, ARF6 is an important regulator of intercellular adhesion^369^ as ARF6 promotes internalization of E-cadherin leading to the disassembly of the adherens junctions^366^. ARF6-GTP can also affect the invasive properties of tumor cells. It has been shown to be an important regulator of invadopodia formation and cell invasion which depends on ARF6-mediated activation of the MEK/ERK signaling pathway^342^. Similar findings were found *in vivo* where sustained activation of ARF6 significantly enhanced the invasive capacity of tumor cells through ARF6-induced ERK signaling and subsequent Rac1 activation to promote invadopodia formation and cell invasion^343^.

Cell migration and invasion occurs through a process of steps. Mesenchymal cells migrate in a five-step process that includes pseudopod formation, focal adhesion contacts with the extracellular matrix, localized proteolysis of the extracellular matrix, actomyosin contraction, and, finally, detachment of the trailing edge of the cell. Epithelial cells, however, migrate and invade using focal adhesion contacts and stress fibers without any proteolysis of the extracellular matrix^370^. To add more complexity, cancer cells can also move in a collective fashion. Pancreatic ductal adenocarcinoma cells have been shown to invade into the surrounding tissue as isolated single cells that leave the primary tumor^221^, small clusters of cells that maintain contact with the primary tumor^318^, or as detached cell clusters that extend into interstitial tissue gaps and along perineural structures^371^.

Single cell invasion and collective cell invasion have the potential to occur at the same time in a primary tumor, however, not in the same exact cell. A cancerous lesion is comprised of a heterogeneous population of tumor cells and tumor microenvironment. For example, in an area of the tumor that is subjected to large amounts of inflammatory stimulus, a cell might be directed to undergo epithelial-to-mesencymal transition and invade individually while other cells in a distant area of the tumor that is not exposed to inflammation may invade collectively. Alternatively, single cell invasion and collective cell invasion may occur in a step-wise pattern. For instance, epithelial-to-mesenchymal transition occurs in a local manner at the invasive front of tumors, and once a single cell detaches from the primary tumor and sets up residence at a secondary site, these cells may then undergo mesenchymal-to-epithelial transition^372^. Depending on the microenvironment at the secondary site, these cancerous cells may then invade collectively. Cell signaling may also determine the method of invasion. TGF-β induces epithelial-to-mesenchymal transition and single cell invasion in breast cancer. Using a rat model of breast carcinogenesis, researchers found that inhibition of TGF-β led to an inhibition of single cell invasion, but saw no interference with collective cell invasion^373^ further suggesting that these two methods of cancer cell invasion can occur in the same tumor at the same time, but not in the same cell.

microRNAs have also been shown to control pancreatic ductal adenocarcinoma invasion. Expression of miR-194, miR-200b, miR-200c, and miR-429 were upregulated in highly metastatic pancreatic ductal adenocarcinoma cell lines while miR-224 and miR-486 have been shown to have significantly higher expression in highly invasive pancreatic ductal adenocarcinoma^374^. Tumor suppressor miRNAs are also dysregulated in pancreatic cancer. For example, miR-146a expression was decreased in metastatic pancreatic cancer cell lines^259^ and ectopic expression of either miR-96 or miR-520h induces inhibition of pancreatic cancer migration and invasion^177,375^. miR-218 also controls cellular invasion. Alajez et al. found that increased levels of miR-218 decreased nasopharyngeal carcinoma cell migration. An increase in miR-218 target Paxillin (due to a decrease in miR-218 expression) leads to increased invasion of non-small cell lung cancers^181^. In gastric cancer, mir-218 expression leads to a decreased in both invasion and metastasis^179^. Finally, in primary pancreatic ductal adenocarcinoma samples, a low level of miR-218 expression correlates with metastatic disease^184^.

To determine if miR-218 plays a role in the inhibition of pancreatic ductal adenocarcinoma invasion, Matrigel-coated Boyden chamber assays were performed. When cells were transfected with the precursor to miR-218, invasion of both cell lines was significantly reduced. Alternatively, transfection using the miR-218 antagomir partially rescued the invasive phenotype. Interestingly, this decrease in invasion was not reliant on the cell’s dependency on KRAS. This suggests that miR-218 is part of a signaling axis that controls pancreatic ductal adenocarcinoma invasion independently from signaling pathways downstream of KRAS. This finding has the possibility of opening up new potential biomarkers for predicting pancreatic cancer metastasis or drug targets to provide novel therapeutics for patients with metastatic disease.

There are three stages in the life of an invadopodium that have been proposed: initiation, assembly, and maturation. Initiation occurs through activation of Src by growth factor receptors leading to the activation of signaling cascades, and, ultimately, actin polymerization^335^. Invadopodia assembly requires localized actin polymerization. Release of cofilin creates severed actin filaments. Subsequent dephosphorylation of cortactin then promotes actin filament elongation creating a branched actin network throughout the pseudopod^336^. Maturation is defined by the ability of an invadopodium to secrete matrix metalloproteinases. Membrane-bound MT1-MMP is delivered to the plasma membrane of invadopodia by three different mechanisms: 1) endocytic recycling, 2) from intracellular stores, and 3) through the actin cytoskeleton and exocytic machinery^335^. Neel et al. determined that the majority of pancreatic cancer cell lines develop invadopodia, and that KRAS activity is both necessary and sufficient for invadopodia formation. Interestingly, the pathway downstream of KRAS responsible for invadopodia formation was the RalB effector pathway, specifically through RalB/RLIP76^340^. RalB is a GTPase that acts as a GTP sensor for GTP-dependent exocytosis of core vesicles. It is required to stabilize the assembly of the exocyst complex and to localize functional exocyst complexes to the leading edge of migrating cells^376^. The function of RalB compares to the function ARF6 has with the exocyst complex which is to specify delivery and insertion of recycling membranes to regions of the plasma membrane undergoing dynamic reorganization through interaction with the exocyst complex^348^. Botta et al. confirmed that activated KRAS increased invadopodia formation in human pancreatic ductal epithelial cells, but only when they were cultured in three dimensions that included a basement membrane analog^355^. β1 integrin was also found to be required for the formation of mature invadopodia in pancreatic cancer cells plated in both two- and three-dimensional conditions^356^. Finally, the guanine nucleotide exchange factor Vav1 has too been shown to be a regulator of matrix degradation in pancreatic cancer by promoting the formation of invadopodia^341^ through activation of Cdc42.

To assess the effect of miR-218 on extracellular matrix degradation by invadopodia, matrix degradation assays were employed. Briefly, sterile coverslips were coated with FITC-conjugated fibronectin. Cancer cells were then plated onto the coverslips for twenty-four hours and stained to visualize phalloidin and DAPI. When cells were transfected with the precursor to miR-218, the area of fibronectin degradation itself and the area of fibronectin degradation per cell number was significantly decreased. On the other hand, when cells were transfected with the miR-218 antagomir, the area of fibronectin degradation itself and the area of fibronectin degradation per cell area was partially rescued in Capan-1 and was fully rescued in Hs 766T. The decrease in extracellular matrix degradation was not contingent on the cell’s dependency on KRAS. This suggests that the pathway miR-218 uses to suppress pancreatic ductal adenocarcinoma invasion regulates invadopodia maturation and the delivery of MT1-MMP to the invadopodia plasma membrane to cause extracellular matrix degradation.

This work suggests that miR-218 is a suppressor of pancreatic ductal adenocarcinoma invasion through a pathway that regulates invadopodia maturation. It is likely that the pathway involved is associated with the delivery of matrix metalloproteinases to the invadopodia plasma membrane (Figure 4.5). Since ARF6 is a novel target of miR-218, I propose the delivery of MT1-MMP may involve pathways connected with protein trafficking. Future directions would be to further elucidate the mechanism behind how miR-218 regulates ARF6-mediated transport of MT1-MMP and to find other potential pathways modulated by miR-218 that are also independent of KRAS.

**Figure 4.5:**
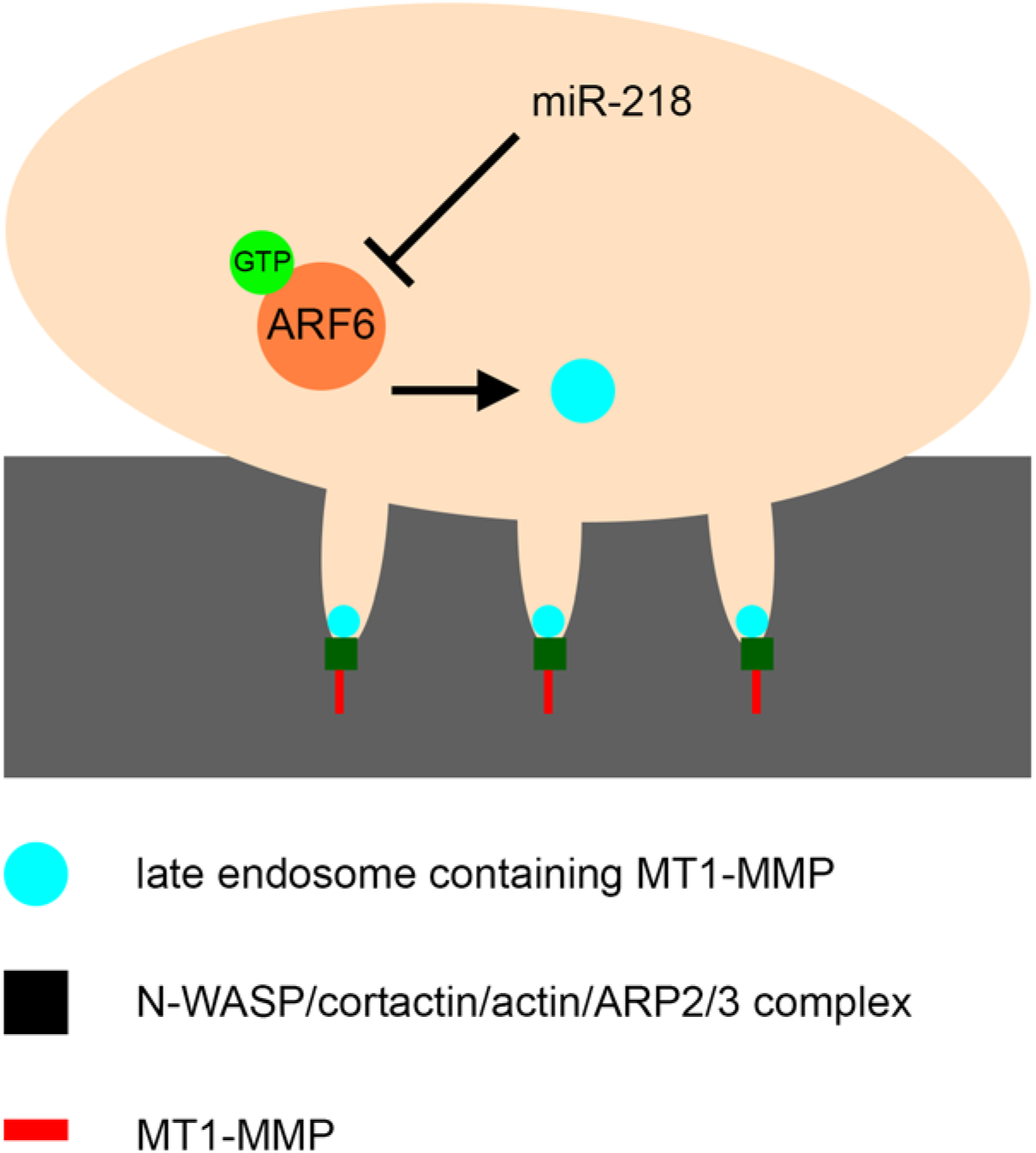
miR-218 is a suppressor of invadopodia maturation. miR-218 inhibits the invasive capability of invadopodia through inhibiting the delivery of MT1-MMP to the plasma membrane of invadopodia by modulation of *ARF6* mRNA expression. miR-218 expression from its putative alternative promoter decreases *ARF6* expression. Depleted *ARF6* levels then prevent trafficking of MT1-MMP-containing late endosomes to the plasma membrane of invadopodia leading to decreased extracellular matrix degradation.

